# scHiCyclePred: a deep learning framework for predicting cell cycle phases from single-cell Hi-C data using multi-scale interaction information

**DOI:** 10.1101/2023.12.12.571388

**Authors:** Yingfu Wu, Zhenqi Shi, Xiangfei Zhou, Pengyu Zhang, Xiuhui Yang, Jun Ding, Hao Wu

## Abstract

While scRNA-seq offers gene expression snapshots, it misses the spatial context of chromatin organization crucial for cell cycle regulation. Single-cell Hi-C, capturing chromatin’s three-dimensional (3D) architecture, fills this void, revealing interactions between genomic regions that transcript-only data might overlook. We introduce scHiCyclePred, a model that utilizes single-cell Hi-C’s multi-scale interaction data to predict cell cycle phases by extracting chromatin’s 3D features. This fusion-prediction model integrates three feature sets into a unified vector. Remarkably, scHiCyclePred outperforms methods like NAGANO and CIRCLET and traditional machine learning techniques across various metrics. Our insights into 3D chromatin dynamics during the cell cycle further underscore its utility. By offering a more comprehensive view of cell cycle dynamics through chromatin structure, scHiCyclePred stands to significantly advance our understanding in cellular biology and holds potential to catalyze breakthroughs in disease research. Access scHiCyclePred at github.com/HaoWuLab-Bioinformatics/scHiCyclePred.

## Introduction

The cell cycle is a highly complex process that involves dynamic changes in various cellular components, including RNA, DNA, and proteins^1–5^. To study the dynamics of the cell cycle, it is essential to analyze the relationship between the cell cycle phases and the state of these cellular components, which form the basis of this fundamental biological process.

Fluorescence imaging is a powerful tool for understanding the relationship between subcellular processes and cellular behavior. Computational analysis of cell-to-cell heterogeneity in single-cell RNA-sequencing data has revealed hidden subpopulations of cells^6–14^. Previous studies have utilized fluorescence imaging to predict cell cycle phases, such as Ersoy et al.’s fast implementation of multi-phase graph partitioning active contours (fastGPAC)^15^, which integrated fluorescently tagged feature distribution with a support vector machine (SVM) to predict cell cycle phases using images of the fluorescently tagged protein GFP-PCNA. Du et al.^16^ developed a cell cycle phases classification algorithm based on 3D fluorescent images of the chromatin marker histone-GFP that extracted 3D intensity, shape, and texture information and combined weighted-SVM and neural network algorithms. Schonenberger et al.^17^ proposed a workflow for classifying cell cycle phases in PCNA-immunolabeled cells using high-quality single-time-point images based on the unique patterns of PCNA distribution. Despite their effectiveness, these approaches are laborious, expensive, and low throughput, posing a significant challenge to research into the dynamic regulation of the cell cycle.

The advent of high-throughput single-cell technologies has significantly elevated the dimensions of single-cell data. This development has provided a novel perspective for deducing cell cycle phases. Among single-cell technologies, single-cell RNA-sequencing (scRNA-seq) data has emerged as an extremely useful tool for examining cellular heterogeneity in gene expression with unprecedented precision. Consequently, it has been widely employed for identifying cell cycle phases. A case in point is Hsiao et al.^18^, who utilized scRNA-seq data analysis to characterize and infer quantitative cell cycle phases. Kowalczyk et al.^19^ employed scRNA-seq data to uncover changes in cell cycle phases during the aging of hematopoietic stem cells.

The emergence of single-cell Hi-C technology has revolutionized the study of chromatin’s three-dimensional structure^20^. The ability to determine cell cycle phases from single-cell Hi-C data is critical for analyzing and comprehending changes in chromatin’s spatial structure during various cell cycle phases. This knowledge is indispensable for revealing cell cycle dynamics. However, accurately predicting cell cycle phases directly from single-cell Hi-C data remains a formidable challenge. Consequently, some studies have focused on constructing the pseudo-trajectory sequence of the cell cycle. For example, Nagano et al.^21^ obtained single-cell Hi-C data from mouse embryonic stem cells (ESCs) at different cell cycle stages and utilized machine learning algorithms to calculate the ‘repli-score’, ratio of short-range connections and frequency of mitotic interactions for each cell. These indicators enabled the cells to follow the pseudo-trajectory of cell cycles. Subsequently, we call this method NAGANO. Ye et al.^22^ proposed a method for constructing cell cycle pseudo-trajectories based on the combination of multiple feature sets called CIRCLET. The study generated four distinct feature sets and their combinations as inputs for CIRCLET, and the pseudo-trajectory sequence of single cells was reconstructed by calculating the distance between cells using the dimensionality reduction method (Wishbone), constructing the K-Nearest Neighbor (KNN) graph between cells based on the calculated distance, and dividing the pseudo-trajectory sequence into two semicircle trajectories using the KNN graph. However, direct prediction of cell cycle phases using only the constructed cell cycle pseudo-trajectory sequence remains challenging, as it is necessary to know the number of cells contained in each cell cycle phase. Suppose that there are *M* cells in the G1 phase, *N* cells in the early-S phase, *P* cells in the mid-S phase, and Q cells in the late-S/G2 phase. The precise implementation is to sort the pseudo-trajectory values corresponding to each cell in ascending order, with the prediction results for the 1 to *M* cells being the G1 phase, the *M* + 1 to *M* + *N* cells being the early-S phase, the *M* + *N* + 1 to *M* + *N* + *P* cells being the mid-S phase, and the *M* + *N* + *P* + 1 to *M* + *N* + *P* + *Q* cells being the late-S/G2 phase. Despite the potential of constructing the pseudo-trajectory sequence of the cell cycle from single-cell Hi-C data, the lack of general datasets containing information on the number of cells in each cell cycle phase renders it impossible to predict the cell cycle phases of individual cells. Moreover, the prediction effect of this method remains unsatisfactory. Therefore, accurate and user-friendly computational methods for predicting cell cycle phases based solely on single-cell Hi-C data are urgently needed.

To overcome the hurdles of predicting cell cycle phases from single-cell Hi-C data, we present a computational framework, scHiCyclePred. This framework integrates multiple feature sets extracted from single-cell Hi-C data and employs a fusion-prediction model based on deep learning methods to predict cell cycle phases. We also propose two feature sets, the bin contact probability feature set, and a small intra-domain contact probability feature set, to improve the accuracy of cell cycle phase prediction. Furthermore, we benchmark the performance of scHiCyclePred against existing methods and demonstrate that it outperforms them in predicting cell cycle phases. Finally, we analyze the changing patterns of chromatin’s three-dimensional structure during the four cell cycle phases using a model interpretation approach. Overall, our proposed framework provides an accurate and user-friendly computational method for predicting cell cycle phases based solely on single-cell Hi-C data and sheds new light on understanding the dynamics of chromatin during the cell cycle.

## Results

### Overview of scHiCyclePred

The deep learning-based framework of scHiCyclePred consists of two crucial steps: the extraction of multiple feature sets and a fusion-prediction model (Figure 1). The former extracts features of chromatin’s three-dimensional structure from the single-cell Hi-C data based on multi-scale interaction information. This step involves extracting the following feature sets: (1) contact probability distribution versus genomic distance feature set from the overall cellular interaction information, (2) bin contact probability feature set from the overall chromatin interaction information, and (3) small intra-domain contact probability feature set from the intra-domain interaction information on chromatin. To integrate the knowledge of multi-scale interactions in cells and intuitively predict the cell cycle stage, we develop a fusion-prediction model that integrates the three feature sets generated by the convolution module.

**Figure 1.**
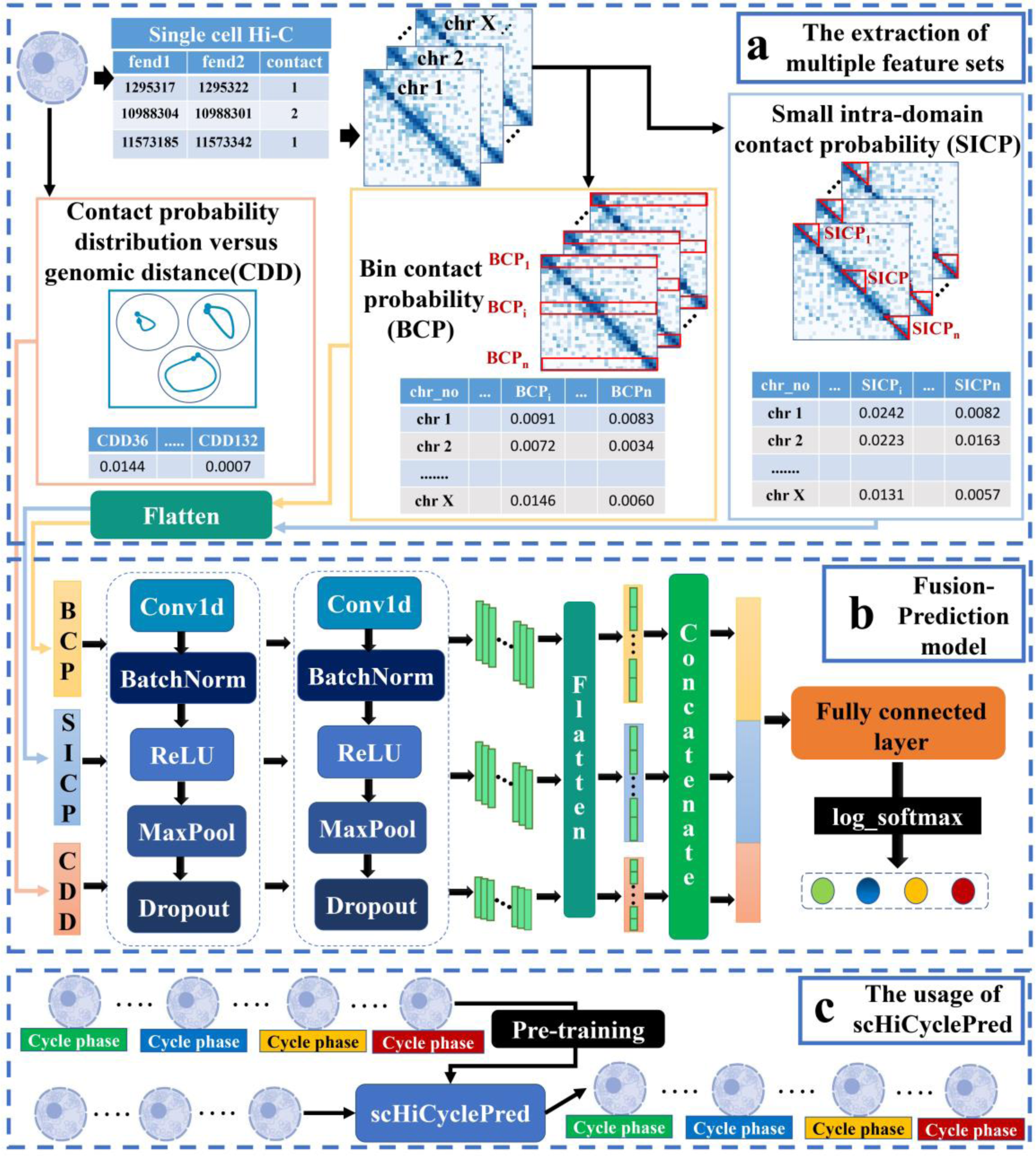
The framework of scHiCyclePred. **(a)** The extraction of multiple feature sets. scHiCyclePred combines read pair locus mapping file and chromatin interaction pair file to generate a unique chromosome contact matrices for each chromosome in every cell and extract three feature sets, contact probability distribution versus genomic distance (CDD), bin contact probability (BCP) and small intra-domain contact probability (SICP). **(b)** Fusion-Prediction model. We develop a deep learning model that combines convolution and feature fusion modules to accurately predict cell cycle phases. **(c)** The usage of scHiCyclePred. The model is initially pre-trained using labeled single-cell Hi-C data. Subsequently, this pre-trained model can be applied to classify new cell data.

In the fusion-prediction model, three feature vectors for each cell are input into the model, which generates three vectors in parallel after passing through two convolution modules composed of a Conv1d layer, BatchNorm layer, Maxpool layer, and Dropout layer, followed by a flattening process. These three generated vectors are then merged into a single vector. The scores from different categories are mapped using a linear layer and “log_softmax”, and the classification outcome is determined by the index with the highest score. In the following sections, we provide a detailed description of the workflows of the two steps in the scHiCyclePred framework.

### Effectiveness evaluation of single feature set and multiple feature sets

To demonstrate the effectiveness of the multi-scale contact probability feature sets, we validate and analyze the performance of our extracted feature sets in this section. To accomplish this, we input each of the three feature sets into the network model independently and validate the classification performance obtained by using each feature set separately. To ensure the robustness of our results, we partition the NAGANO dataset into fifty independent training and testing sets by altering the random seeds, with the NAGANO dataset being the total of each training and testing set. We then evaluate the prediction performance of the three feature sets using four evaluation metrics: accuracy (ACC), F1 Score (F1), AUC, and AP. Specifically, the AUC value and AP value represent the area under the ROC curve and the area under the Precision-recall (PR) curve, respectively (Figure 2A).

**Figure 2.**
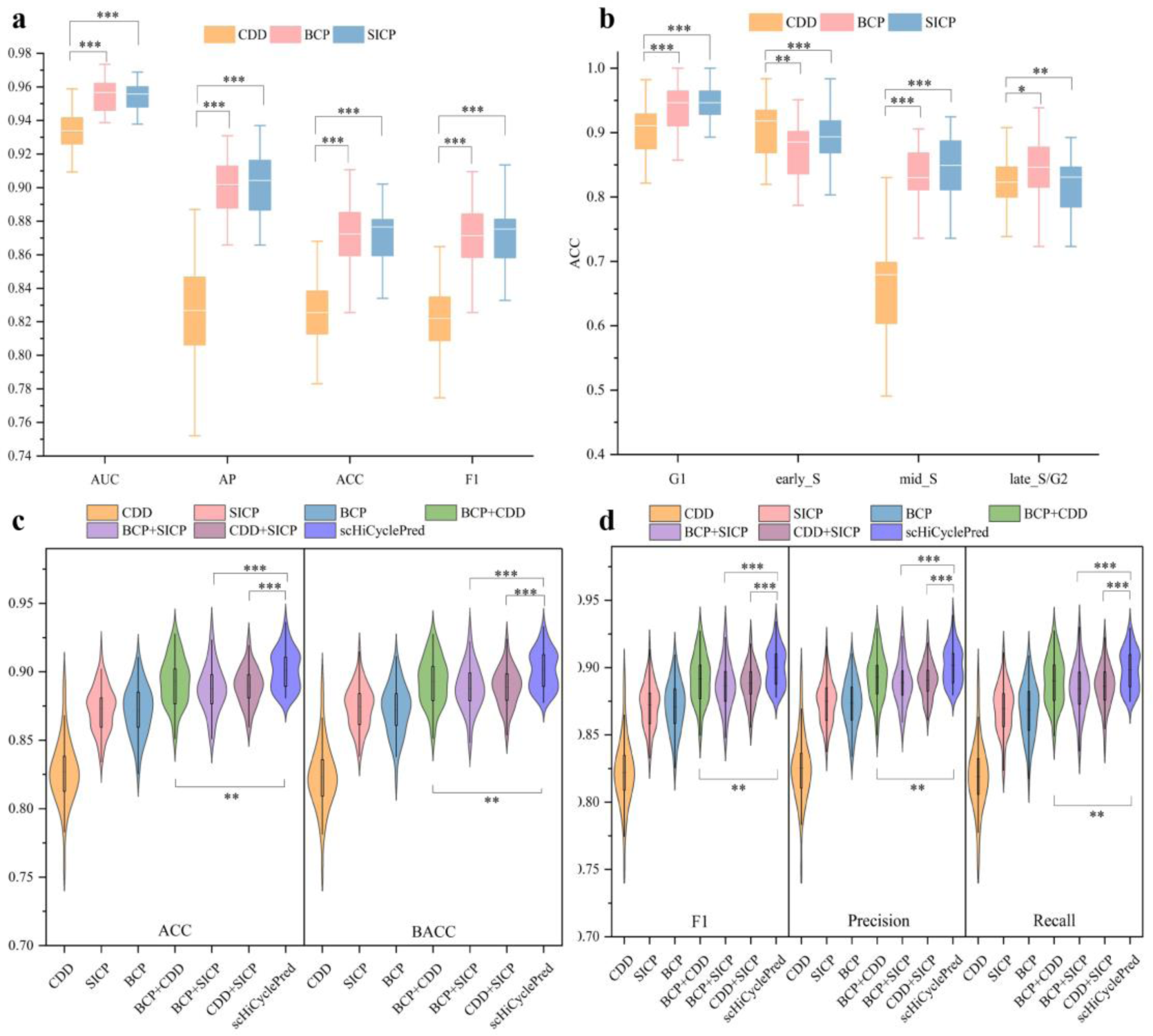
The effectiveness evaluation of the single feature set and the multiple feature sets. **(a)** The performance evaluation of three single feature sets. **(b)** The performance evaluation of three single feature sets according to the cell cycle phases, where the lower and upper edges represent the minimum value and the maximum value of the result, respectively. The bottom edge of the box represents the first quartile (Q1) and the top edge of the box represents the third quartile (Q3). The white line represents the median value. **(c)** and **(d)** The performance comparison of single feature set and the multiple feature sets, where the low end and top of vertical line represent the lower adjacent value and the upper adjacent value of the result, respectively. The bottom edge of the box represents the first quartile (Q1) and the top edge of the box represents the third quartile (Q3). The black line represents the median value. The larger the area of the graph of a region, the greater the probability of a distribution around a certain value. (*** indicates P-value < 1×10^-3^ , ** indicates P-value < 1×10^-2^, * indicates P-value < 5×10^-2^).

The results indicate that the three feature sets are effective for predicting the four cell cycle phases (Figure 2B). The detailed prediction results of the three feature sets for different phases show that: (1) In the G1, early_S, and late_S/G2 phases, the results of the three feature sets are similar; (2) In the mid-S phase, the results of the SICP and BCP feature sets are significantly larger than those of the CDD feature set (Figure 2B).

To evaluate the effectiveness of our fusion-prediction model based on multiple feature sets, we compare the performance of the three-feature-set fusion model with that of six other models constructed by retaining the corresponding feature extraction modules: three single-feature-set models (CDD, BCP, and SICP) and three two-feature-set fusion models (BCP-CDD, CDD-SICP, and BCP-SICP). We utilize the fifty independent training and testing sets mentioned in the previous section for this experiment. We use four evaluation metrics, namely ACC, F1, Precision, and BACC^23–25^, to assess the prediction performance of each model (Figure 2C and Figure 2D).

The effectiveness of our extracted features is demonstrated by the fact that the three single-feature-set models yield higher performance (Figure 2C and Figure 2D). Furthermore, feature fusion enhances the performance of the three two-feature-set models, highlighting the importance of our feature fusion. The final results indicate that the fusion-prediction model with three feature sets exhibited the best performance in terms of ACC, F1, Precision, and BACC, with the least difference between the maximum and minimum values, i.e., the most stable result. Based on these comparison results, our fusion-prediction model can effectively fuse feature sets at all scales and deliver superior cell cycle phase prediction ability.

### Performance evaluation of scHiCyclePred in predicting cell cycle phases

To demonstrate the superiority of our proposed scHiCyclePred method, we compare it to two cell cycle trajectory construction methods (NAGANO and CIRCLET) using four evaluation metrics in this section. Although cell cycle trajectory construction is not primarily designed for predicting cell cycle phases, in order to evaluate NAGANO’s and CIRCLET’s results efficiently, we generate predictive labels based on cell number and cycle order. As the NAGANO and CIRCLET methods do not require the division of training and testing sets, both methods performed predictions directly on the original NAGANO dataset. To compare scHiCyclePred with these two methods, we use them to calculate the average of the ACC, F1, and Precision metrics obtained from fifty testing sets. Additionally, we evaluate the effectiveness of our constructed deep learning model by employing three conventional machine learning methods, namely Support Vector Machine^26^, Logistic Regression^27^, and Random Forest^28,29^. The same fifty independent training and testing sets mentioned earlier are utilized for this experiment (Figure 3).

**Figure 3.**
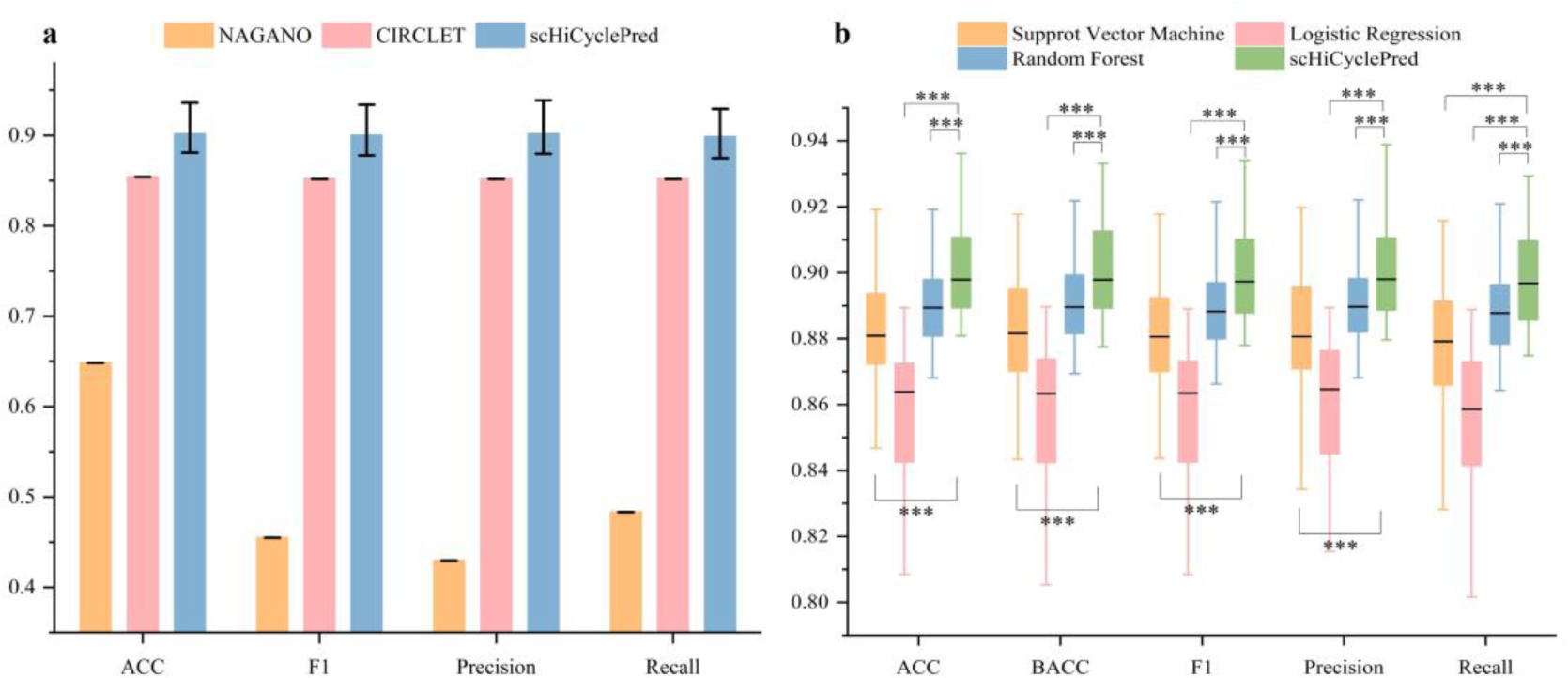
The performance comparison of scHiCyclePred with other methods. **(a)** The performance comparison of scHiCyclePred with NAGANO and CIRCLET. The top of the bar graph represents the average value, the bottom of the vertical line. **(b)** The performance comparison of scHiCyclePred with other machine learning methods. The lower and upper edges represent the minimum value and the maximum value of the results, respectively. The bottom edge of the box represents the first quartile (Q1), the top edge of the box represents the third quartile (Q3), and the white line represents the median value. (*** indicates P-value < 1×10^-3^).

The results indicate that scHiCyclePred outperforms NAGANO and CIRCLET in three metrics: ACC, F1, and Precision (Figure 3A). Furthermore, scHiCyclePred still achieves optimal performance in terms of ACC, F1, Precision, and BACC metrics compared to using the three conventional machine learning methods (Figure 3B). These findings demonstrate that the scHiCyclePred method achieves superior performance in predicting cell cycle phases.

### Robustness validation of scHiCyclePred

We evaluate the scHiCyclePred model in two separate approaches to validate its robustness on datasets for the following purposes: (1) Validating the effectiveness of scHiCyclePred on imbalanced datasets by downsampling the original NAGANO dataset. (2) Testing the effectiveness of scHiCyclePred on the newly drop-processed datasets. For the downsampling experiment, we downsample the NAGANO dataset into nine imbalanced datasets (A-I) composed of different proportions (Figure 4A). Using a ratio of 8:2, the nine datasets are split into two parts: a training set and a testing set. The training set is used to train the model, while the testing set is used to evaluate its performance. The comparison results of the four evaluation metrics of ACC, F1, Precision, and BACC demonstrate the effectiveness of scHiCyclePred on imbalanced datasets (Figure 4C and Figure 4D).

**Figure 4.**
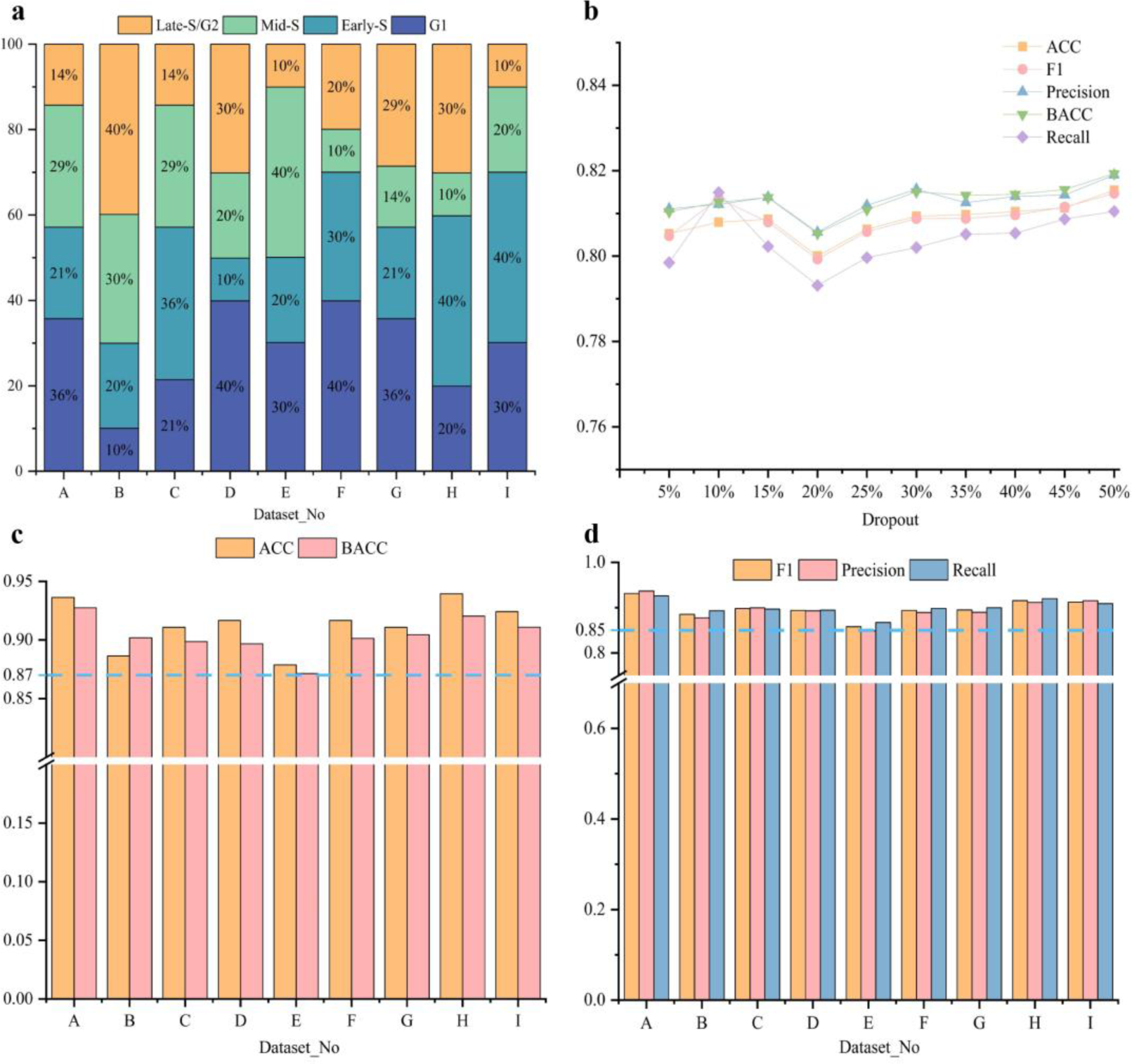
The performance of scHiCyclePred on imbalanced datasets and the newly drop-processed datasets. **(a)** The number of cells at each cell cycle phase in different datasets. **(b)** The performance of scHiCyclePred on the newly drop-processed datasets. **(c)** and **(d)** The performance of scHiCyclePred on imbalanced datasets.

For the drop experiment, the NAGANO dataset’s chromatin interaction pair file (raw_data file) is split into five separate training sets (raw_data_train file) and testing sets (raw_data_test file). Each NAGANO dataset is the sum of each corresponding training and testing set. Then, for each set of raw_data_test files, 5%-50% (5%, 10%, 15%, 20%, 25%, 30%, 35%, 40%, 45%, 50%) rows of data are randomly selected, where y is the total number of rows in the raw_data_test file. Taking into account that interaction pairs with contact numbers of 1 or 2 account for 98.8% of the total number of interaction pairs (details regarding the number of interaction pairs with varying contact numbers, as well as their corresponding proportions) (Supplementary Figure S1), we modify the number of contacts between two segments (counting columns) for the selected rows using the following procedure: adding or subtracting one at random from the number of contacts. Following this procedure, each training set is matched to 10 testing sets, resulting in a total of 50 testing sets across the five training sets. Each group of experiments is trained using its corresponding training set, and only the 10 testing sets in the same group as the current training set are utilized to test the trained model to prevent data leakage. The comparison results of ACC, F1, Precision, and BACC metrics demonstrate the effectiveness of scHiCyclePred on the newly drop-processed datasets (Figure 4B).

Overall, scHiCyclePred performs well in predicting imbalanced datasets on all four evaluation metrics, namely ACC, BACC, F1, and Precision (Figure 4C and Figure 4D). These results suggest that scHiCyclePred is robust on various imbalanced datasets. Moreover, the scHiCyclePred prediction performance on the testing sets obtained using the drop strategy does not decline noticeably with an increase in the drop rate. Instead, the performance essentially stabilized. These results indicate that scHiCyclePred has strong robustness on drop-processed datasets. In summary, the results from both experiments demonstrate the effectiveness and robustness of scHiCyclePred in predicting cell cycle phases on various datasets.

### Analysis of chromatin change patterns across various cell cycle phases

We conduct an analysis to further investigate the pattern of changes in the 3D structure of chromatin during different cell cycle phases. Specifically, we utilize the SHapley Additive exPlanations (SHAP) method^26,28^ to analyze the feature importance of the three feature sets during the four cycles. The SHAP graph displays the vertical axis representing features and the dots representing samples, with redder colors indicating higher feature values and bluer colors indicating lower feature values. The horizontal axis demonstrates the SHAP value, where positive values indicate a positive effect on the prediction, and negative values indicate a negative effect.

Regarding the CDD feature set, we focus our analysis on the top 20 significant features. The results of the CDD feature set’s importance evaluation illustrate that 12 features are present among the top 20 features of the four cell cycle phases: CDD_61-70_, CDD_72_, and CDD_74_ (Figure 5). Confirming previous findings^22^, the proportion of short-distance chromatin interaction pairs increases gradually with the cell cycle phases based on the contact distance and SHAP value represented by these features. In addition, CDD_111_, CDD_117_, and CDD_119_ appear in the 20 most important features of early-S and late-S cycles, indicating that there may be more significant changes in long-distance contacts from early-S to late-S/G2 cycles.

**Figure 5.**
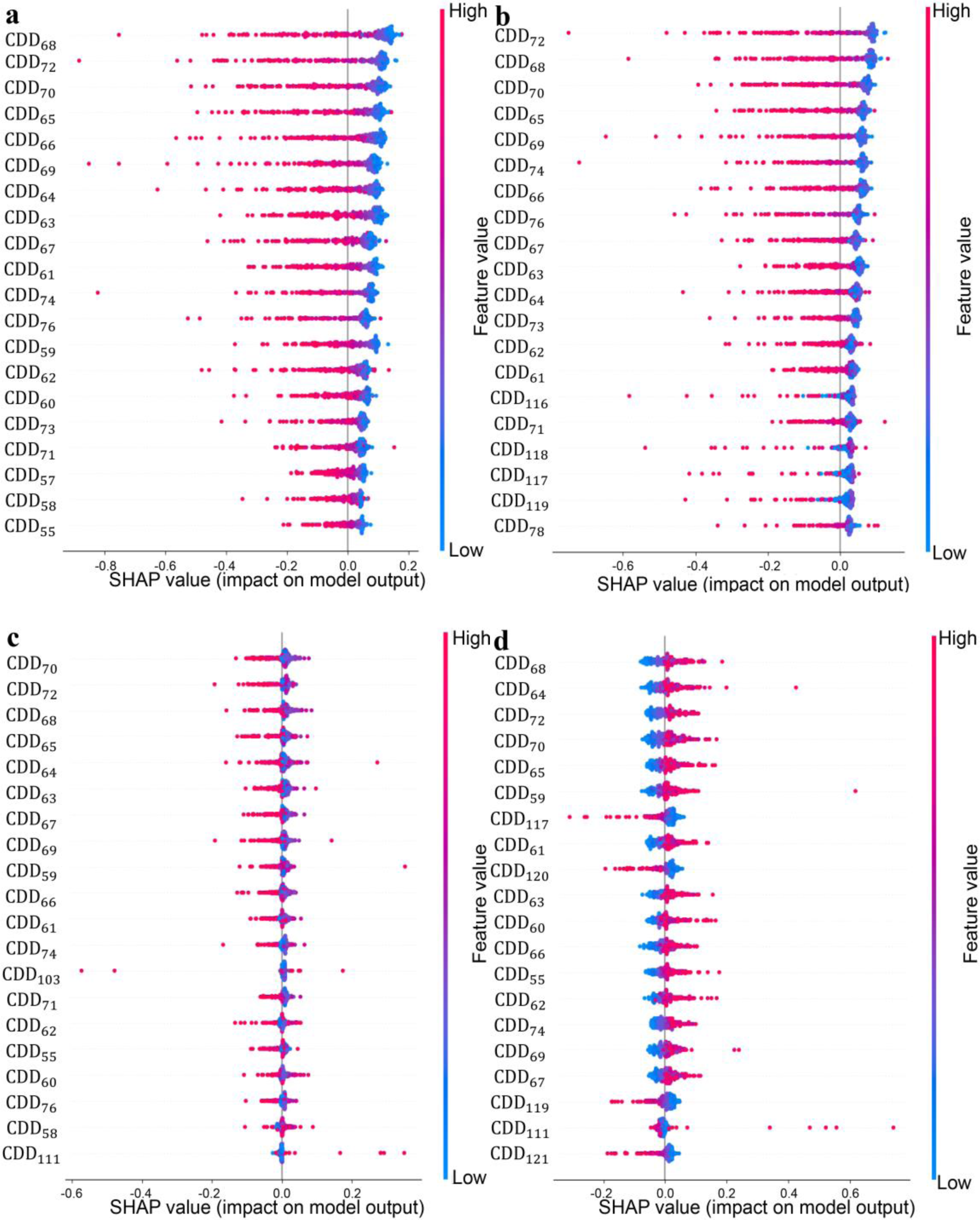
The importance evaluation results of the first 20 features in the CDD feature set. **(a)** G1 phase. **(b)** Early-S phase. **(c)** Mid-S phase. **(d)** Late-S/G2 phase. We analyze the top 20 significant features of the CDD feature set and find that 12 features are present across all four cell cycle phases: CDD_61-70_, CDD_72_, and CDD_74_.

For the BCP feature set and SICP feature set, we primarily analyze the top 50 important features. The results of the BCP feature set’s importance evaluation demonstrate that for fragments on different chromatin, their three-dimensional structure change patterns vary during the cell cycle phases, such as BCP_41_ of chr15 (its three-dimensional structure gradually loosens) and BCP_98_ of chr6 (its three-dimensional structure gradually tightens), BCP_64_ of chr4 (gradual loosening), and BCP_86_ of chr16 (gradual tightening (Figure 6)). In addition, the feature importance results for bins from the same chromatin reveal that the change patterns of the three-dimensional structure of the majority of adjacent bins are relatively similar in cell cycle phases.

**Figure 6.**
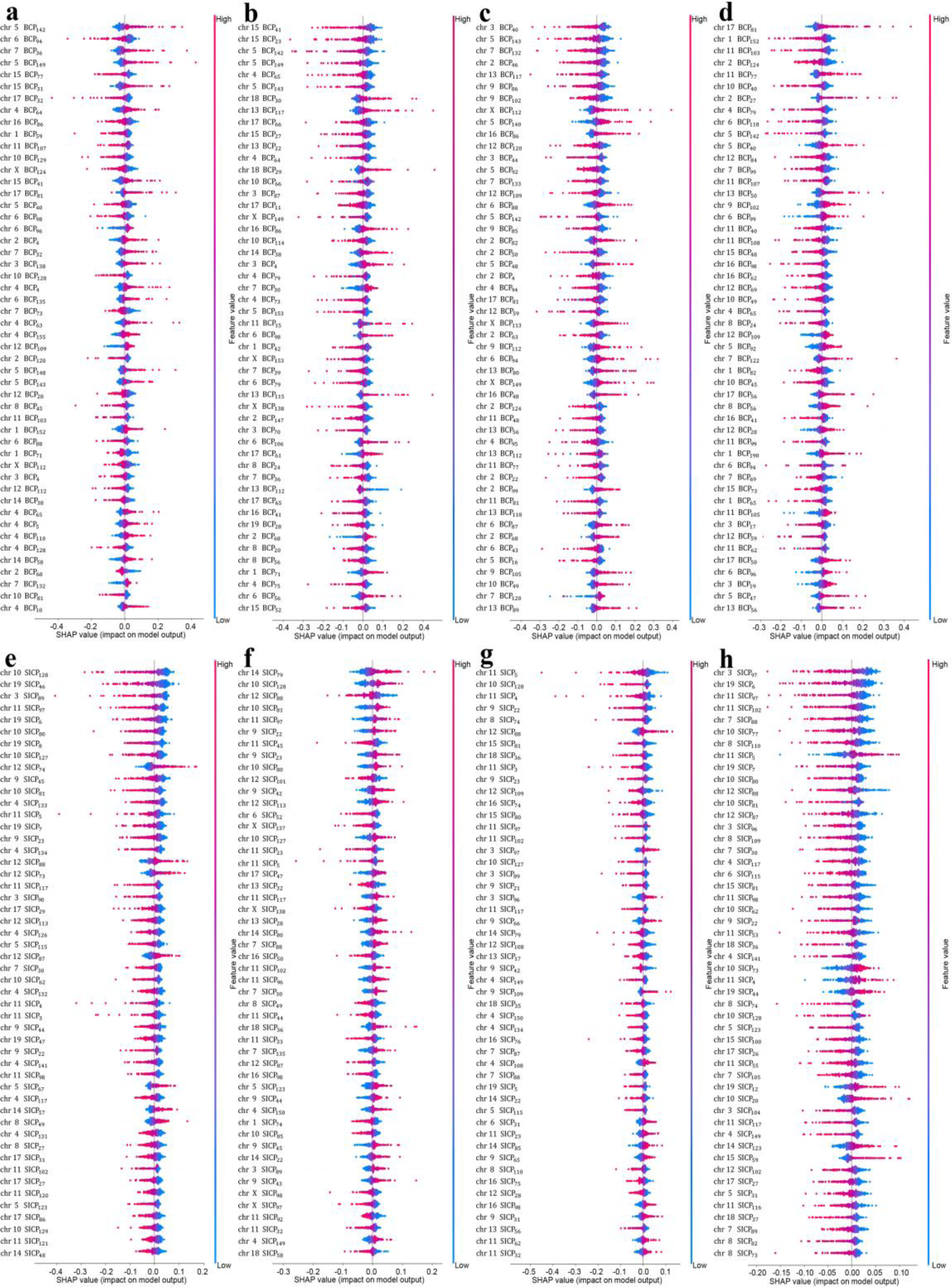
The importance evaluation results of the first 50 features in the BCP and SICP feature set. **(a)** BCP feature set in G1 phase. **(b)** BCP feature set in Early-S phase. **(c)** BCP feature set in Mid-S phase. **(d)** BCP feature set in Late-S/G2 phase. **(e)** SICP feature set in G1 phase. **(f)** SICP feature set in Early-S phase. **(g)** SICP feature set in Mid-S phase. **(h)** SICP BCP feature set in Late-S/G2 phase.

The results of the SICP feature set’s importance evaluation show that: (1) For G1 and Lata-S/G2 cycles, the most prominent three-dimensional structure of the bin neighborhoods tends to be loose; (2) The three-dimensional structure change patterns of adjacent bins are similar, such as SICP_127_ and SICP_128_ of chr10, SICP_3-5_ of chr11, SICP_87_ and SICP_88_ of chr12, etc; (3) From the perspective of the three-dimensional structural changes in the bin neighborhood, the prominent contact patterns of the mid-S cycle and the late-S/G2 cycle are relatively similar (Figure 6). One possible reason for (3) may be that the degree of three-dimensional structure change of chromatin gradually decreases from the mid-S cycle to the late-S/G2 cycle. This finding is consistent with previous study^21^.

In our study, on the basis of feature importance ranking, we meticulously annotate the patterns associated with each stage of the cell cycle and link them to specific genes. To validate the accuracy of these patterns, we conduct an extensive analysis of relevant research literature pertaining to the cell cycle. Our findings are as follows: (1) In the G1 phase, we discover a significant correlation between the gene lists from papers^30–32^ and the genes annotated by our patterns, with the same p-value of 0.005858. (2) During the early-S phase, we observe a strong correlation between the gene lists in papers^33–35^ and the genes annotated through our patterns, with p-values of 2.388×10^-5^, 0.0003518, and 0.001689. (3) Similarly, for the mid-S phase, there is a notable correlation between the gene lists from papers^36–38^ and our annotated genes, with p-values of 9.535×10^-5^, 0.00224, and 0.003723. (4) Finally, in the late-S/G2 phases, we find a substantial correlation between the gene lists in papers^39–41^ and our annotated genes, with p-values of 4.067×10^-6^, 0.0001014, and 0.0006912. In summary, our analysis consistently demonstrates a strong association between our annotated patterns and the cell cycle, reaffirming the precision and relevance of our approach.

## Discussion

In this study, we introduce a new method called scHiCyclePred for predicting cell cycle phases using single-cell Hi-C data. Our method involves the extraction of multiple feature sets based on single-cell Hi-C data and the development of a fusion-prediction model for cell cycle phase prediction. Three feature sets are extracted, including the existing CDD feature set (overall cellular interaction information) and two novel feature sets: the BCP feature set (overall chromatin interaction information) and the SICP feature set (intra-domain interaction information). Our fusion-prediction model successfully combines the three feature sets and improves prediction performance. Furthermore, we evaluate the fusion-prediction model using ablation experiments and compare the performance of scHiCyclePred with other popular methods, including NAGANO, CIRCLET, LR, SVM, and RF, in predicting cell cycle phases. Our results demonstrate that the scHiCyclePred method has good performance in predicting cell cycle phases and is more robust than existing single-cell Hi-C data-based cell cycle phase prediction methods.

In addition, we analyze the pattern of changes in the three-dimensional structure of chromatin during cell cycle phases by evaluating the impact of important features. However, we note that current single-cell Hi-C data is biased due to the coverage consistency of current single-cell Hi-C techniques, which hinders the unraveling of the relationship between cell cycle dynamics and three-dimensional structural patterns of chromatin. Therefore, in the future, it will be necessary to address these biases present in single-cell Hi-C data and further incorporate our method for cell cycle phase prediction. Overall, our study provides a promising approach for predicting cell cycle phases using single-cell Hi-C data, and further research in this field is needed to fully realize the potential of this method. The scHiCyclePred method offers an accurate and user-friendly computational approach to predict cell cycle phases based solely on single-cell Hi-C data, and it provides new insights into understanding the dynamics of chromatin during the cell cycle.

## Materials and methods

### Data preparation

The single-cell Hi-C data used in this study are obtained from the study by Nagano et al.^12^, which includes Hi-C data from 1171 mouse embryonic stem cells labeled according to their cell cycle phase using fluorescence-activated cell sorting (FACS). The cells are classified into four phases: 280 cells in the G1 phase, 303 cells in the early-S phase, 262 cells in the mid-S phase, and 326 cells in the late-S or G2 phase. Raw data files are downloaded from https://github.com/tanaylab/schic2, containing chromatin interaction pair files and read pair locus mapping files. The chromatin interaction pair file contains information such as the sequence number of chromatin fragment pairs and the count of interactions, while the read pair locus mapping file contains the relationship between chromatin fragment sequence number, chromatin information, and precise position information.

Most current computational methods for single-cell Hi-C data represent the data as multiple chromatin contact matrices with a given resolution^42–50^. However, generating a chromatin contact matrix requires specific information about interacting chromatin fragments in each cell, which is not contained in the raw data files. To address this limitation, we combine the read pair locus mapping file and chromatin interaction pair file to generate a unique chromatin interaction information file for each chromatin in each cell. Next, we divide each chromatin in the cell into multiple segments called bins based on a specified resolution *R*. This allows us to represent the single-cell Hi-C data as a matrix of bin-pair interactions, which can be used as input for our proposed scHiCyclePred framework. Finally, we generate the chromatin contact matrix *C* by mapping the interaction information from the chromatin interaction information file to the corresponding bin. However, since Nagano et al. could not capture interaction information for chromatin Y, we generate 20 chromatin contact matrices (*C*_1_,*C*_2_,…,*C*_19_,*C_X_*) per cell to cover all possible combinations of chromatin interactions. This allows us to effectively capture the chromatin interaction patterns and generate accurate representations of the single-cell Hi-C data for use in our proposed scHiCyclePred framework.

## The extraction of multiple feature sets

### Contact probability distribution versus genomic distance feature set

Nagano et al. discovered that the contact probability based on linear distance division showed different states in various cell cycle phases, and Ye et al. proposed the contact probability distribution versus genomic distance feature set to determine the cell cycle trajectory^21,22^. Therefore, we employ the contact probability distribution versus genomic distance (CDD) as a feature set in our framework. It is important to note that the CDD feature set is extracted from the cell-wide interaction information rather than the chromatin contact matrix. In this feature set, the interaction pairs are allocated into intervals based on linear distance, with the linear distance range represented by each interval gradually increasing. The mapping formula for the interaction pairs is as follows:

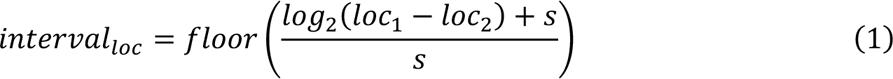

where *loc*_1_ and *loc*_2_ represent the positions of two interacting fragments on the chromatin, and *s* denotes the exponential step length of each interval. Based on the study by Nagano et al. and Ye et al.^22,23^, we set *s* = 0.125 and the distance range for genes to be [2K, 9.3M]. After assigning all contacts, the contact probability distribution versus genomic distance feature set is extracted by calculating the probability of the contact count in each interval as shown in Equation 2.

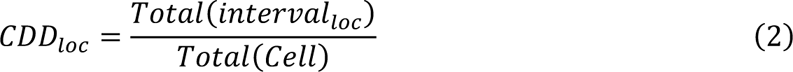

where *Total*(*interval*) represents the contact count in the corresponding bin, and *Total*(*Cell*) represents the total number of chromatin contacts in a cell.

### Bin contact probability feature set

The contact frequency distribution of the same site on the same chromatin varies in different cell cycle phases^51–53^. Therefore, we use the bin contact probability (BCP) set of the chromatin as the contact feature set for chromatin in our framework. Specifically, we use the BCP values in the chromatin contact matrix, rather than the CDD feature set, to extract this set of features. We then use the contact feature set of all chromatin in the cell as one of the extracted feature sets. The chromatin contact matrix is *C*_*n*∗*n*_, where *n* is the number of bins on the chromatin. The bin contact probability of the chromatin is calculated as follows:

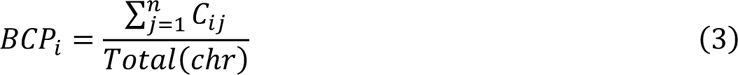

where *Total*(*chr*) represents the total contact count of chromatin at the location of bins and the value range of *i* is [1, n]. The feature set of bin contact probability for each cell is generated by splicing the set of bin contact probability for all chromatin.

### Small intra-domain contact probability feature set

Nagano et al. found that the insulation score of the chromatin domain (INS) is affected by cell cycle dynamics^21^. INS shows the depletion of chromatin interaction information in a domain centered on the target bin. Motivated by this finding, we extract the contact probability of small intra-domain (SICP) on the chromatin as one of the feature sets in our framework. The small domain is defined as the region centered on the target bin and bounded by the adjacent first-order linear bins. Specifically, we calculate the SICP values for each bin by dividing the number of contacts within the small domain by the total number of contacts in the bin. This results in a vector that represents the SICP feature set for the chromatin bin. On the chromosome contact matrix *C*_*n*∗*n*_, the contact probability of a small intra-domain is calculated similarly to the bin contact probability as follows:

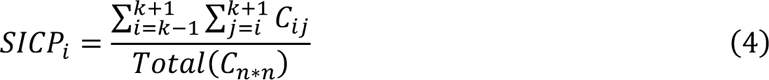

where *C*_k∗k_ represents the currently calculated chromatin domain center and the value range of k is [1, n]. It is important to note that in order to calculate the SICP features, it is necessary to fill matrix *C* with 0 elements, transforming it into matrix *B* . This enables the formation of a complete small domain, even when *k* is equal to 1 and *n*. Similar to BCP, we integrate multiple sets of small intra-domain contact probabilities for entire chromatin as the SICP feature set of a mouse cell.

### Fusion-Prediction model

In this section, we present a deep learning-based fusion-prediction model to accurately predict cell cycle phases by incorporating convolution and feature fusion modules into the architecture. The convolution module generates identical network models for the three feature sets CDD, BCP, and SICP. This module consists of a CNN layer, batch norm layer, max pooling layer, and dropout layer which are stacked twice to generate more complicated features. The CNN layer uses a one-dimensional convolution kernel with a kernel size of 7 and a channel size of 32 to collect features from various input feature sets. The batch norm layer prevents gradient explosion and disappearance, while the max pooling layer reduces feature dimension, preserves key features, scales back model calculations, avoids overfitting, and enhances generalizability. We use ReLU as the activation function to connect the batch norm layer and max pooling layer^54^, adding nonlinear components to enhance the model’s expression capability. The dropout layer effectively prevents model overfitting by discarding some neurons during forward propagation with a predetermined probability. Finally, a flattening operation is applied to the data produced from the second dropout layer to combine the data from all channels into a vector.

In the feature fusion module, the three vectors corresponding to the three feature sets generated by the convolution module are combined into a single vector. The scores from various categories are then mapped using a linear layer, followed by the “log_softmax” function^55–57^. The ultimate classification outcome corresponds to the index with the highest score. Using the “log_softmax” function avoids the value overflow problem and facilitates the calculation of the loss function. To address the issue of imbalanced samples and enhance the overall performance of the model, we adopt the focal loss function^58,59^ as the loss function of the model. The focal loss is calculated as in Equation 5,

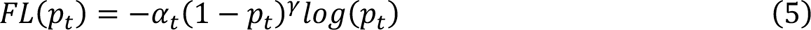

where the value range of γ is [0,5], *p*_*t*_ means the probability that the model predicts the current sample as phase *t*, and −*log*(*p*_*t*_) is utilized to calculate the cross entropy loss. If the *p*_*t*_ corresponding to the current sample phase is smaller, it means that the prediction result of the model is more inaccurate, then the coefficient (1 − *p*_*t*_)^*γ*^ of the difficult sample will increase, and the difficult sample will lose. *⍺*_*t*_ represents the weight coefficient corresponding to phase *t* , and the value is the number of cells contained in phase *t*. This model is trained by minimizing the focal loss.

Additionally, to prevent overfitting of the model to the training set during model building, we employ an early stopping^60–64^ mechanism and a 5-fold cross-validation approach. The model training is terminated if the loss of the model on the validation set does not reduce for 10 consecutive epochs (Figure 1 and Supplementary Figure S2).

## Acknowledgments

We thank members of the group for their valuable discussions and comments. The scientific calculations in this paper have been done on the HPC Cloud Platform of Shandong University.

## Author contributions

These authors contributed equally to this work.

## Data and materials availability

The source code for scHiCyclePred is freely available on GitHub at https://github.com/ HaoWuLab-Bioinformatics/scHiCyclePred.

