## Supplementary Figs. S1-S2 for "scHiCyclePred: a deep learning framework for predicting cell cycle phases from single-cell Hi-C data using multi-scale interaction information"

**Figures:**

Figure S1. Details regarding the number of interaction pairs with varying contact numbers, as well as their corresponding proportions.

Figure S2. The details of the fusion-prediction model.

†These authors contributed equally to this work.

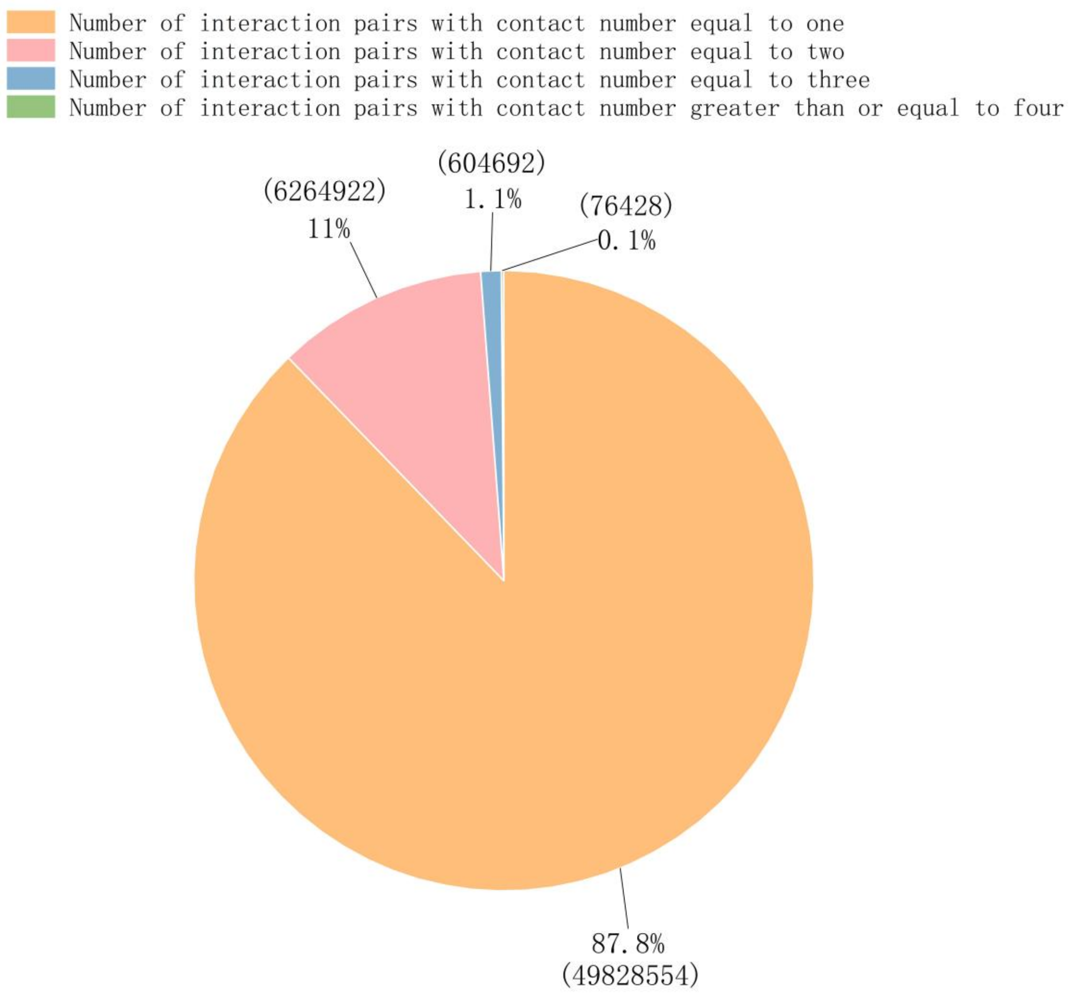


**Figure S1. Details regarding the number of interaction pairs with varying contact numbers, as well as their corresponding proportions.** Interaction pairs with contact numbers of 1 account for 98.8% of the total number of interaction pairs. Interaction pairs with contact numbers of 2 account for 11% of the total number of interaction pairs. Interaction pairs with contact numbers of 3 account for 1.1% of the total number of interaction pairs. The number of interaction pairs with contact numbers greater than or equal to 4 accounts for only 0.1% of the total number of interaction pairs.

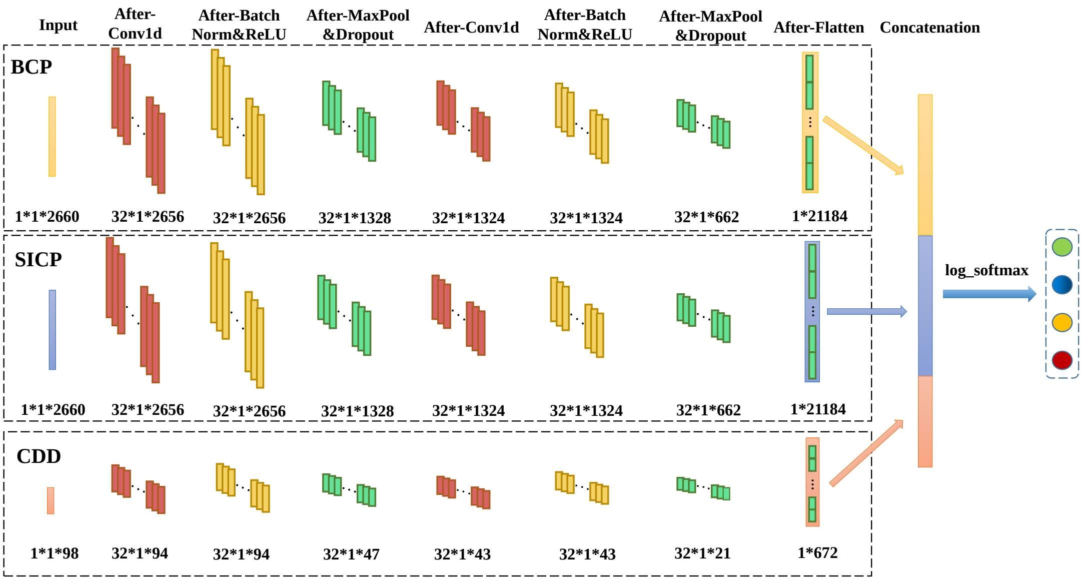


**Figure S2. The details of the fusion-prediction model.**
